# The ZKIR Assay, a novel Real-Time PCR Method for the Detection of *Klebsiella pneumoniae* and Closely Related Species in Environmental Samples

**DOI:** 10.1101/855668

**Authors:** Elodie Barbier, Carla Rodrigues, Geraldine Depret, Virginie Passet, Laurent Gal, Pascal Piveteau, Sylvain Brisse

**Affiliations:** Agroécologie, AgroSup Dijon, CNRS, INRA, Univ. Bourgogne Franche-Comté, Dijon, France; Institut Pasteur, Biodiversity and Epidemiology of Bacterial Pathogens, Paris, France

**Keywords:** *Klebsiella*, phylogroup, soil, detection, screening, ZKIR qPCR, culture method

## Abstract

*Klebsiella pneumoniae* (Kp) is of growing public health concern due to the emergence of strains that are multidrug-resistant, virulent, or both. Taxonomically, Kp includes seven phylogroups, with Kp1 (*K. pneumoniae sensu stricto*) being medically prominent. Kp can be present in environmental sources such as soils and vegetation, which could act as reservoirs of animal and human infections. However, the current lack of screening methods to detect Kp in complex matrices limits research on Kp ecology. Here we analysed 4222 genome sequences and found that existing molecular detection targets lack specificity for Kp. A novel real-time PCR method, the ZKIR assay, was developed and used to detect Kp in 96 environmental samples. Results were compared to a culture-based method using SCAI agar medium coupled to MALDI-TOF mass spectrometry identification. Whole-genome sequencing of environmental Kp was performed. The ZKIR assay was positive for the 48 tested Kp reference strains, whereas 88 non-Kp strains were negative. The limit of detection of Kp in spiked soil microcosms was 1.5 × 10^-1^ CFU g^-1^ after enrichment for 24 h in LB supplemented with ampicillin, and 1.5 × 10^3^ to 1.5 × 10^4^ CFU g^-1^ directly after soil DNA extraction. The ZKIR assay was more sensitive than the culture method. Kp was detected in 43% of environmental samples. Genomic analysis of the isolates revealed a predominance of phylogroups Kp1 (65%) and Kp3 (32%), a high genetic diversity (23 MLST sequence types), a quasi-absence of antibiotic resistance or virulence genes, and a high frequency (50%) of O-antigen type 3. This study shows that the ZKIR assay is an accurate, specific and sensitive novel method to detect the presence of Kp in complex matrices, and indicates that Kp isolates from environmental samples differ from clinical isolates.

**IMPORTANCE:** The *Klebsiella pneumoniae* species complex (Kp) includes human and animal pathogens, some of which are emerging as hypervirulent and/or antibiotic resistant strains. These pathogens are diverse and classified into seven phylogroups, which may differ in their reservoirs and epidemiology. Proper management of this public health hazard requires a better understanding of Kp ecology and routes of transmission to humans. So far, detection of these microorganisms in complex matrices such as food or the environment has been difficult due to a lack of accurate and sensitive methods. Here, we describe a novel method based on real-time PCR, which enables detection of all Kp phylogroups with high sensitivity and specificity. We used this method to detect Kp isolates from environmental samples, and show based on genomic sequencing that they differ in antimicrobial resistance and virulence gene content, from human clinical Kp isolates. The ZKIR PCR assay will enable rapid screening of multiple samples for Kp presence and will thereby facilitate tracking the dispersal patterns of these pathogenic strains across environmental, food, animal and human sources.

## INTRODUCTION

*Klebsiella pneumoniae* (Kp) is one of the leading causes of multidrug resistant (MDR) healthcare-acquired infections, with increasing rates of resistance to carbapenems and other last resort antibiotics being reported (1, 2). Furthermore, Kp is also an important agent of severe community-acquired infections (so-called ‘hypervirulent’ strains) in healthy persons (3), with recent worrisome reports of convergence between hypervirulent and MDR phenotypes (1, 4). Kp is recognized as a colonizer of the throat and the intestinal tract of humans and animals (5–7).

The main sources of human exposure to Kp are not well defined. Previous studies highlighted the large distribution of Kp in outdoor environments, including water, sewage, soil and plants (8–13). Animal and human food, particularly retail meat or salad may also be contaminated (6, 14, 15). Many studies suggest that food, water and/or environmental exposure may be associated with virulent and/or antibiotic resistant Kp to humans (13, 14, 16, 17). However, little is known on the relative contribution of these different sources of transmission. Although such information is a prerequisite to control efficiently transmission routes and reduce exposure, the ecology of Kp is currently poorly understood.

The systematics of Kp has been refined through recent taxonomic updates, which highlighted the existence of seven phylogroups (Kp1 to Kp7), corresponding to distinct taxa, within *Klebsiella pneumoniae sensu lato*. The *K. pneumoniae* species complex includes 5 different species: *K. pneumoniae sensu stricto* (phylogroup Kp1), *K. quasipneumoniae* [subsp. *quasipneumoniae* (Kp2) and subsp. *similipneumoniae* (Kp4)], *K. variicola* [subsp. *variicola* (Kp3) and subsp. *tropica* (Kp5)], ‘*K. quasivariicola’* (Kp6) and *K. africana* (Kp7) (8, 18–20). Most of these taxa are still often misidentified as “*K. pneumoniae”* or *“K. variicola”* due to the unsuitability of traditional clinical microbiology methods to distinguish among members of the Kp complex. Henceforth, we will use the ‘Kp’ abbreviation to refer collectively to the seven taxa of the *K. pneumoniae* species complex, and will reserve ‘*K. pneumoniae*’ for *K. pneumoniae sensu stricto* (*i.e.,* phylogroup Kp1). As all Kp are potentially pathogenic for humans and animals and can share acquired resistance and virulence genes, it is important that the seven taxa be considered together when investigating the routes of transmission and ecology of Kp.

Detection of Kp is not well integrated in food or environmental microbiological surveillance programs and there is a general lack of tools and procedures for its detection and quantification. Culture-based laboratory methods used for the detection of microorganisms in complex matrices are time consuming and have a low throughput. Moreover, Kp culture methods have not been validated so far for food safety screening. Some molecular methods (without need of sequencing) have been proposed over the years for the rapid detection of Kp (21–25). They target the 16S-23S rRNA internal transcribed spacer sequence (ITS) (21), coding sequences of *tyrB* (25), *khe* (24, 26), chromosomal beta-lactamase (*bla*) genes (27, 28) or other molecular targets (22). Some of these targets are described as able to detect the Kp complex (21), while others were designed for specific members of the Kp complex, such as Kp1 and Kp3 (22, 23).

Real-time PCR is a powerful approach for the rapid detection and quantification of microorganisms in complex matrices (29). This approach presents multiple advantages including easy standardization and high throughput. The aims of this work were (i) To define the phylogenetic distribution of previously proposed molecular targets for Kp detection, in light of recent taxonomic updates; and (ii) To develop a real-time PCR method for the rapid, specific and sensitive detection of all Kp members. (iii) In addition, we used the novel qPCR method to detect Kp in environmental samples and explored the genomic features (including antibiotic resistance and virulence genes) of the recovered Kp isolates.

## MATERIALS AND METHODS

### Bacterial strains and culture conditions

A panel of 136 strains from the collections of the Institut Pasteur, Paris (France) and INRA Dijon (France) was used (Supplementary material Table S1). It included the *K. pneumoniae* type strain ATCC13883^T^ and 47 other strains of the Kp complex (Kp1 to Kp7 phylogroups, including type strains) isolated from patients, animals and outdoor environments from multiple geographic locations. Eighty-eight non-Kp bacterial strains from other species of the genus *Klebsiella*, closely related genera (*Raoultella, Enterobacter*), and from other species that have similar environmental niches were included for comparison. All strains were regenerated by streaking on tryptone soya agar (TSA, Conda, Spain). After 24h at 37°C, single colonies were transferred to 10 ml tryptone soya broth (TSB, Conda, Spain). Bacterial suspensions were collected after 24 h incubation at 37°C for further experiments.

### Bacterial DNA extraction

The bacterial DNA was extracted according to a homemade protocol. One ml of the 24h bacterial culture was centrifuged (7000 g, 10 min, 4°C), the pellet washed with sterile water and suspended in 800 µl Tris (50 mM, pH 8) in which 10 µl EDTA (500 mM, pH 8), 115 µl lysozyme (10 mg/ml, incubation 37°C, 30 min), 57 µl SDS 20% (incubation 65°C, 15 min), 5 µl RNAse (10 mg/ml, incubation 37°C, 30 min) and 23 µl proteinase K (6 mg/ml, incubation 37°C, 1h30) were added. Finally, a 1:1 volume of chloroform/ isoamyl alcohol (25/24/1) was added. The suspension was shaken 3 min at 15 rpm, then centrifuged 3 min at 20 000 g. The supernatant was transferred in a new tube and a 1:1 volume of chloroform/ isoamyl alcohol (24/1) was added. After shaking (3 min, 25 rpm) and centrifugation (3 min, 20 000 g), this step was repeated once. After transfer of the supernatant in a new tube, a 1:10 volume of NaCl (5 M) was added (shaking 1 min, 15 rpm). Finally, a 1:1 volume of isopropanol (shaking 3 min, 15 rpm) was added. Precipitated DNA filaments were collected with a glass Pasteur pipette and transferred in 400 µl frozen ethanol (70%). After shaking 3 min at 15 rpm, DNA filaments were collected, dried at room temperature and finally dissolved in 100 µl TE (pH 8). DNA concentration was estimated using NanoDrop 2000 Spectrophotometer (ThermoFischer Scientific, USA) and adjusted at 20 ng/µl. Purified DNA was stored at −20°C.

A simpler protocol was implemented for the PCR specificity assay and validation study. Bacteria grown in TSB or enrichment medium were centrifuged 5 min at 13 000 g. Pellets were suspended in 200 µl of Tris HCl (5 mM pH 8.2) supplemented with 13 µl of Proteinase K (1 mg/ml). After 2 h incubation at 55 °C, Proteinase K was inactivated during 10 min boiling. Cell debris were discarded by centrifugation (5 min at 13 000 g) and the supernatant was stored at – 20°C before use for real-time PCR.

### Soil DNA extraction

Soil metagenomic DNA was extracted from 2 g aliquots of soil using the modified ISO standard 11063 method (30). Crude soil DNA extracts were first purified onto PVPP (polyvinylpolypyrrolidone) minicolumns (BIORAD, France), then with the GeneClean Turbo kit (MP Biomedicals, France) according to the manufacturer’s protocol (30). Purified DNA concentrations were determined using a Nanodrop 2000 spectrophotometer.

### *In silico* identification of the target DNA region

The phylogenetic distribution of molecular targets that were previously described in the literature for Kp detection, were investigated *in silico*. All available *Klebsiella* genomes were downloaded from the NCBI database (February 2018) and combined with the ones of Institut Pasteur’s internal reference collection (n=66) (20, 31, 32), representing a total of 4,222 genomes which included *Klebsiella* species and closely related *Raoultella* species. The average nucleotide identity (ANI) metric was used to classify the genomes in each of the phylogroups. To avoid redundancy in the dataset, unique representatives of each MLST sequence type (ST) were selected for the analysis, leading to 1,001 unique genomes. Nucleotide BLAST (BLASTN) was used to detect the presence of the target sequences in the 1,001 genomes, with 80% nucleotide identity and 80% length coverage as cutoffs. Primer-blast (https://www.ncbi.nlm.nih.gov/tools/primer-blast/) was used (with default parameters) to check the distribution of primer target sites in the genomic sequences. Specificity was defined as the proportion of target organisms among those expected to be amplified by the target assay. Sensitivity was defined as the proportion of target organisms expected to be amplified by the target assay. A phylogenetic tree was constructed (33) and the output was visualized using iTOL (https://itol.embl.de/; Figure S1).

### Design of primers

The Kp *tyrB* and *khe* gene sequences (accession numbers: AF074934.1 and AF293352.1) were aligned against the genome sequences of *K. pneumoniae* (CP012744.1, CP012743.1, CP028787.1), *K. variicola* (CP008700.1, CP013985.1, CP017289.1), *K. quasipneumoniae* (CP014071.1, CP023478.1, CP029432.1) and *K. quasivariicola* (CP022823.1) using BLASTN to identify conserved genomic regions within the Kp complex. Sequences of closely related bacterial species *K. oxytoca* (CP027426.1), *Raoultella ornithinolytica* (CP010557.1), *R. planticola* (CP026047.1) and *K. aerogenes* (CP014029.2) were added in the alignments to confirm the specificity of these regions for the Kp complex.

Primer sets were designed using Primer3Plus (http://www.bioinformatics.nl/cgi-bin/primer3plus/primer3plus.cgi). The specificity of the predicted primer sets and amplicons was checked by applying BLASTN on the GenBank nucleotide collection (nr/nt) from the NCBI database and Multiple primer analyzer (https://www.thermofisher.com) was run to check for dimer formation. The best candidate primer sets defined by this *in silico* approach were ordered for synthesis at Eurogentec; among these was primer pair ZKIR_F (5’-CTA-AAA-CCG-CCA-TGT-CCG-ATT-TAA-3’) and ZKIR_R (5’-TTC-CGA-AAA-TGA-GAC-ACT-TCA-GA– 3’).

### Real-time PCR

All real-time PCR assays were performed on an ABI StepOne™ Real-time thermocycler (Fisher Scientific, France) with the following temperature program: 95 °C for 3 min and 40 cycles at 95 °C 10 s and 60 °C for 1 min. Melt curves were generated with temperature increments of 0.3 °C per cycle from 60 to 95 °C. DNA was amplified in a 20 µl PCR mix containing 10 µl of Takyon Low Rox SYBR MasterMix dTTP Blue (Eurogentec, Belgium), 2 µl of each primer (final concentration 300 nM), 2.5 µl of template DNA and 3.5 µl PCR grade water. For detection of Kp from environmental samples, 0.5 µl of T4 Gene 32 Protein (Sigma Aldrich) was added to the PCR mix. The specificity and cross-reactivity of the ZKIR assay was evaluated with 2 ng of purified DNA of 48 Kp complex strains and 88 non-Kp isolates (Table S1).

Two universal primers targeting a 174 base pairs region of the *16S rRNA* gene of Eubacteria, 341f (5’-CCT-ACG-GGA-GGC-AGC-AG-3’) and 515r (5’-ATT-CCG-CGG-CTG-GCA-3’), were used for positive control PCR reactions as described in a previous report (34).

### Standard curve development and sensitivity assessment

Aliquots of 2.5 µl adjusted to 7.5 ng, 750 pg, 75 pg, 7.5 pg, 750 fg, 375 fg, 45 fg and 15 fg of genomic DNA of *K. pneumoniae* ATCC13883^T^ were prepared in triplicate and amplified with the optimized PCR conditions described above. Results were analysed with the StepOne™ Data Analysis software. PCR efficiency was determined. The genome number expressed as logarithm was plotted against Ct values and the correlation coefficient (R^2^) of the standard curve was calculated.

### Comparison of the ZKIR PCR and culture methods for detection of Kp in environmental samples

Ninety-six environmental samples collected between July and September 2018 (23 in July; 39 in August and 34 in September) in Auxonne (Burgundy, France) were analysed for Kp presence using the ZKIR assay and culture methods in parallel. Samples corresponded to bulk soils (n=32), roots (31), leaves (29) and irrigation water (4) and were processed in the lab within 24 hours after sampling. Ten g of soil were weighed in 180 ml pots (Dutscher, France). Plant leaves and roots were properly cut, cleaned with sterile water and transferred in 180 ml pots. Processed samples were suspended in 90 ml of lysogeny broth (LB, 5 g of NaCl, 5 g of yeast extract and 10 g of Tryptone for a 1 l final volume) supplemented with ampicillin (10 mg/l, ampicillin sodium salt, Sigma Aldrich). Five hundred millilitres of water samples were filtered through a 0.25 µm membrane (Millipore, France). The membrane was incubated in 20 ml of LB supplemented with ampicillin as described above. After 24 h of incubation at 30 °C, enrichments were vortexed and 500 µl aliquots were centrifuged (5 min at 5800 × g) and washed with sterile water. The pellet was suspended in 500 µl of sterile water and boiled for 10 min. Boiled enrichments were ten-fold diluted (1:10 and 1:100) and the dilutions were used as template for real-time PCR.

In parallel, enrichments were serially diluted (1:10 to 1:10 000) in sterile water before plating on Simmons Citrate Agar enriched with inositol (SCAI medium) (35). Plates were incubated 48 h at 37 °C. Each plate was screened for presumptive colonies of *Klebsiella* (large, yellow, dome-shaped colonies) and ten candidate colonies were purified on SCAI medium and identified using Matrix Assisted Laser Desorption Ionization Time of Flight (MALDI-TOF) Mass Spectrometry (MALDI Biotyper, Bruker) according to the MALDI Biotyper Compass database version 4.1.80 (Bruker Daltonics, Germany). In addition, whole-genome sequencing (WGS) was performed for environmental Kp isolates (n=31) using the NextSeq-500 sequencing platform (Nextera XT library; 2×150 nt). Genomic assemblies were obtained using SPAdes v3.9. Multilocus sequence typing (MLST) and core-genome MLST (cgMLST) were performed using the BIGSdb-Kp web tool (https://bigsdb.pasteur.fr/klebsiella/klebsiella.html). This resource coupled with Kleborate (https://github.com/katholt/Kleborate) was used to search for antibiotic resistance, virulence and heavy metal tolerance genes, and to predict capsular types. PlasmidFinder was used to look for plasmid replicons (https://cge.cbs.dtu.dk/services/PlasmidFinder/). To construct the tree based on *nif* cluster genes (**Figure S2**), the DNA region between *nifQ* and *nifJ* genes was extracted from the *nif* carrying strains and from a set of reference strains carrying this cluster (36). Sequences were aligned with MUSCLE, and IQ-TREE (http://iqtree.cibiv.univie.ac.at) was used to reconstruct the maximum likelihood phylogeny using the HKY+I+F+G4 model.

### Determination of the limit of detection of the ZKIR assay in artificially spiked soils

Two soils with contrasted edaphic characteristics (Sandy soil A; clay soil V) and free of indigenous Kp, according to the above procedure, were used. A bacterial suspension of *K. pneumoniae* ATCC13883^T^ was serially diluted in sterile water and dilutions were enumerated on TSA plates. Five grams aliquots of each soil were spiked with these decreasing Kp dilutions. Soil samples were prepared in triplicates with three independently grown inoculums at 4 dilutions (36 spiked microcosms for each soil). Negative controls were prepared by adding the same volume of sterile water (3 microcosms for each soil). All 78 spiked and control soil microcosms were enriched for 24 h at 30 °C in 45 ml of LB supplemented with ampicillin (10 mg/l, Sigma Aldrich). Five hundred µl aliquots of enrichment broth were sampled before and after the 24-h enrichment step, centrifuged 5 min at 5800 g and washed with sterile water. The pellet was suspended in 500 µl of sterile water and boiled for 10 min. Boiled enrichments were serially ten-fold diluted (1:10 to 1:100 000) and the diluted suspensions were used for real-time PCR. Moreover, metagenomic DNA was extracted from spiked soils as described above in ‘Soil DNA extraction’ paragraph.

### Data availability

The detailed ZKIR qPCR operating procedure was made publicly accessible to the scientific community through the protocols.io platform (https://dx.doi.org/10.17504/protocols.io.7n6hmhe). Genomic sequences generated in this study were submitted to the European Nucleotide Archive and are accessible under the BioProject number PRJEB34643.

## RESULTS

### Revisiting the phylogenetic distribution of proposed molecular targets for Kp detection

Fourteen molecular targets were found in the literature (Figure S1; Table S2)(21–27, 37). Four of them were proposed to detect the Kp complex, while others were designed for specific members of this complex (*K. pneumoniae*, *K. variicola* or *K. quasipneumoniae;* Table S2). Mapping of the presence/absence of the sequence region expected to be amplified by the primers was performed across *Klebsiella* phylogenetic diversity (**Figure S1**). Regarding the targets proposed for the entire Kp complex, this *in-silico* analysis revealed that only *khe* and *tyrB* (24, 25) presented both a high sensitivity and a high specificity. We further analysed *in silico* the primers target sites for these two genes using primer-blast. The results showed that *khe* primers would be expected to amplify the same region in *Raoultella* spp. (77 bp, with a sequence identity of the homologous region < 80%, hence not visible on Figure S1), as well as an additional region (348 bp) in these organisms. Furthermore, *tyrB* primers also appear to be able to amplify the target region in *K. aerogenes* and *Raoultella* spp., consistent with the distribution of the target region (**Figure S1**). Regarding the targets that were proposed to be specific for Kp1, target KpI50233a (23) proved to be the most specific and sensitive one (although with ability to also amplify *K. aerogenes* isolates), whereas the ones proposed for Kp3 (22) appeared unspecific or to lack sensitivity in our *in silico* analysis (**Figure S1**). Regarding the targeted chromosomal class A beta-lactamases - *bla*_SHV,_ *bla*_OKP_ and *bla*_LEN_ (27), there appeared to be a lack of specificity (**Figure S1),** but this is explained by the high degree of sequence identity between these *bla* genes, even though they represent distinct targets. *In silico, bla*_OKP_ and *bla*_LEN_ primers were specific for Kp2/Kp4 and Kp3/Kp5, respectively. However, proposed *bla*_SHV_ primers (27) appeared to lack specificity and/or sensitivity and also amplified taxa outside the Kp complex (such as *E. coli* or *Salmonella* spp.)

### ZKIR primers design, PCR assay development and optimization

The *tyrB* and *khe* genes were previously proposed as targets for the specific detection of Kp (24, 38). As shown above, *tyrB* and *khe* targets were not totally specific for the Kp complex. We therefore attempted to define primers within the coding sequence of these genes but external to the previously proposed target fragments. However, high identity was observed with non-Kp complex species when blasting the entire coding sequence (data not shown). Consequently, Kp complex-specific primers could not be designed within the coding sequence of these two genes. Interestingly, investigation of sequences upstream of *khe* did reveal a sequence that was highly conserved within the Kp complex. This region located in the intergenic region (IR) upstream of *khe* is partly deleted in other species such as *K. oxytoca* and *Raoultella* spp. As this 249 bp non-coding IR is located between *zur* (zinc uptake regulator) and *khe* (annotated as a putative hemolysin) genes (Figure 1), it was named ZKIR for *zur-khe* intergenic region.

**Figure 1.**
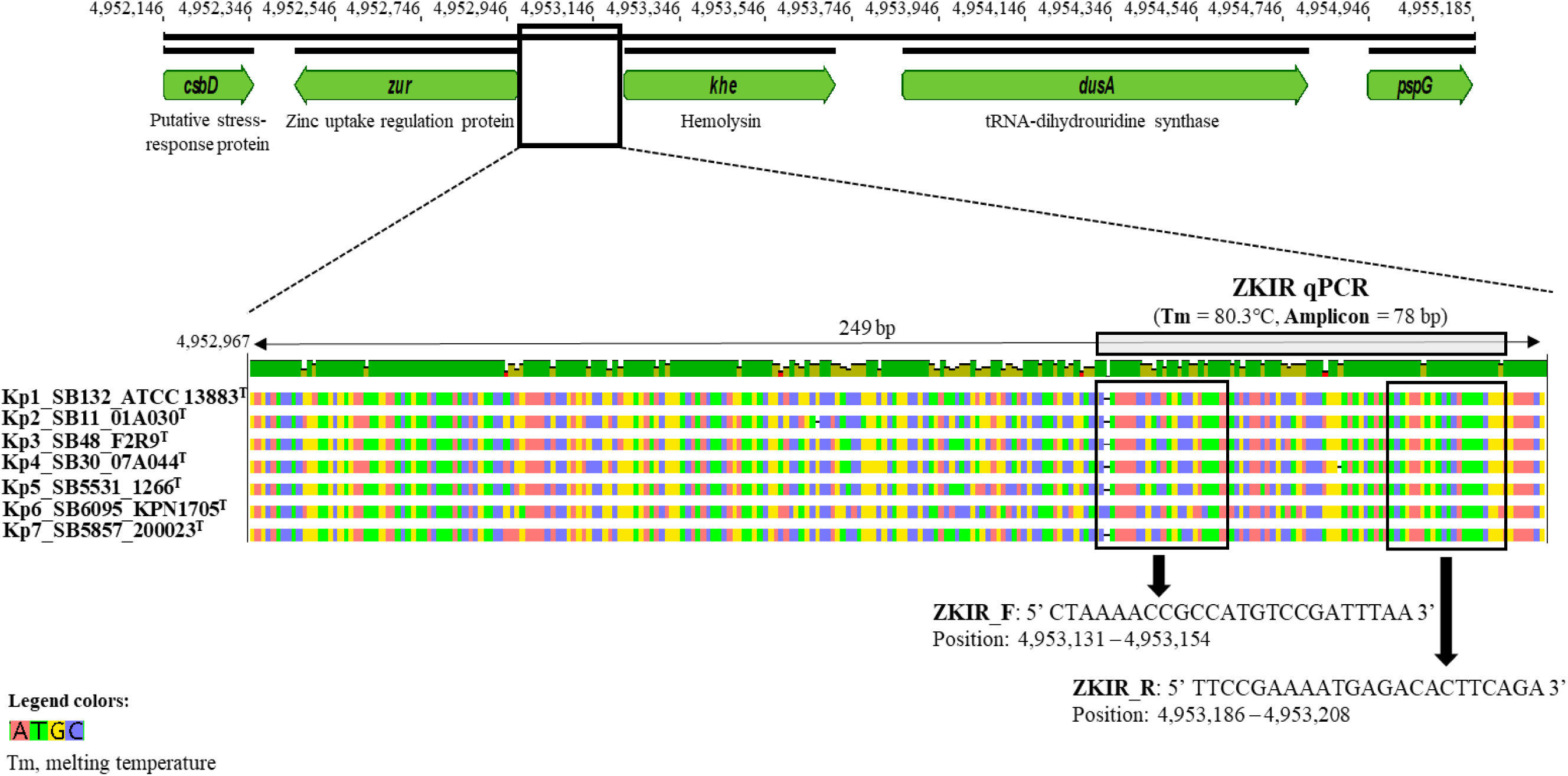
Genetic context of ZKIR region on the genome of strain *K. pneumoniae* ATCC 13883^T^ (GenBank accession number GCA_000742135.1) and detailed location of ZKIR primers and amplicon within the 249-base pair (bp) region that is specific for the *K. pneumoniae* species complex (boxed area).

A pair of primers (ZKIR_F and ZKIR_R, Figure 1) targeting the ZKIR region was designed and tested *in silico.* When implemented in the SYBR green PCR assay using reference strain ATCC 700603 (known as *K. pneumoniae*, but belonging in fact to *K. quasipneumoniae* subsp. *similipneumoniae*), these primers successfully amplified a 78 bp sequence, with a melting temperature of 80.3 °C. Sequencing of PCR products confirmed amplification of the target sequence.

The sensitivity and specificity of the primers were experimentally tested on 2 ng of purified DNA of representatives of Kp phylogroups Kp1 to Kp7 (Table 1) and non-Kp strains. The ZKIR_F and ZKIR_R primers amplified the target sequence in all tested isolates of the Kp complex, with a Threshold Cycle (Ct) value ranging from 13 to 26 and melting temperatures of 80.1 to 80.7 °C. In contrast, when the ZKIR PCR was performed on 88 non-Kp isolates, no amplification was observed, showing that the PCR did not yield false positives. Late, non-specific amplification was recorded with few isolates in the last cycles of the reaction (after cycle 35) and melting temperatures were clearly lower or higher than 80°C, indicating a non-specific amplification. All assays performed on non-Kp samples can therefore be considered negative.

**Table 1.** Comparison between ZKIR qPCR and culture results obtained using samples collected in Auxonne, France, between July and September 2018.

| Source | No. of samples | qPCR positive samples (%) | Culture positive samples (%) |
| --- | --- | --- | --- |
| **Bulk soil** | 32 | 13 (40.6%) | 13 (40.6%) |
| **Roots** | 31 | 19 (61.3%) | 16 (51.6%) |
| **Leaves** | 29 | 8 (27.6%) | 7 (24.1%) |
| **Water** | 4 | 1 (25.0%) | 1 (25.0%) |
| **Total** | 96 | 41/96 (42.7 %) | 37/96 (38.5%) |

### Analytical sensitivity of the ZKIR PCR

PCR reactions were performed with quantities of genomic DNA of *K. pneumoniae* ATCC13883^T^ ranging from 7.5 ng (2.5 × 10^6^ genomes) to 15 fg (5 genomes) (Figure 2 A and B). Linearity of the assay was evaluated by plotting the Ct values against the log_10_ calculated genome number (calculated genome size is about 3.0 fg of DNA per cell, based on a 5.54 Mb *K. pneumoniae* genome). Ct values were closely proportional to the logarithm of the genome number (R^2^=0.99) (Figure 3). The sensitivity of the assay was determined as 15 genomes (45 fg) per reaction mix. No amplification was observed at the lower DNA concentration of 15 fg corresponding to 5 genomes.

**Figure 2.**
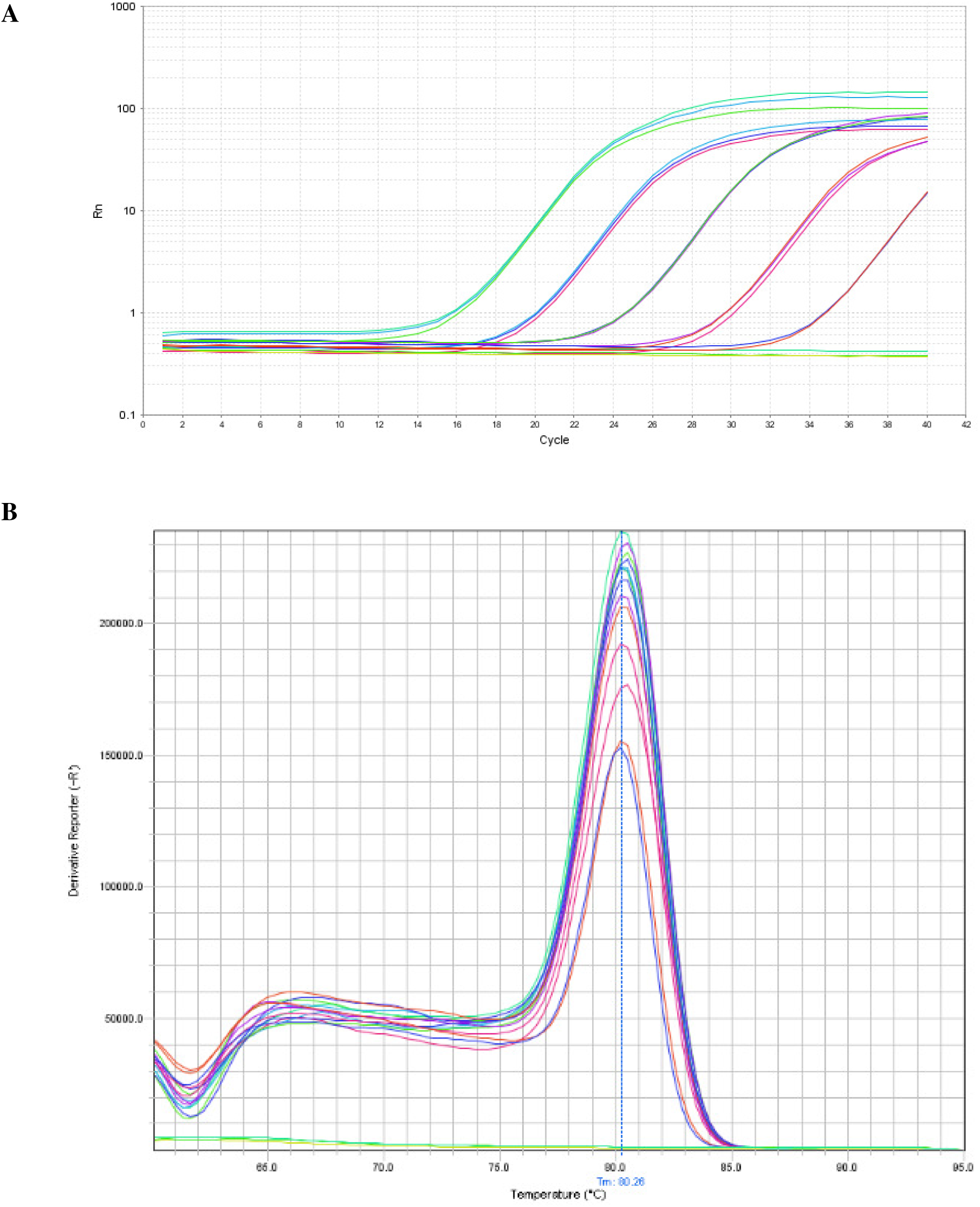
Amplification curves (A) and melt curve peaks (B) established using real-time PCR targeting the ZKIR region with serial dilutions of *K. pneumoniae* ATCC13883^T^ DNA. Triplicates of DNA concentrations of 7.5 ng, 750 pg, 75 pg, 7.5 pg and 750 fg are presented, whereas lower dilutions are not.

**Figure 3.**
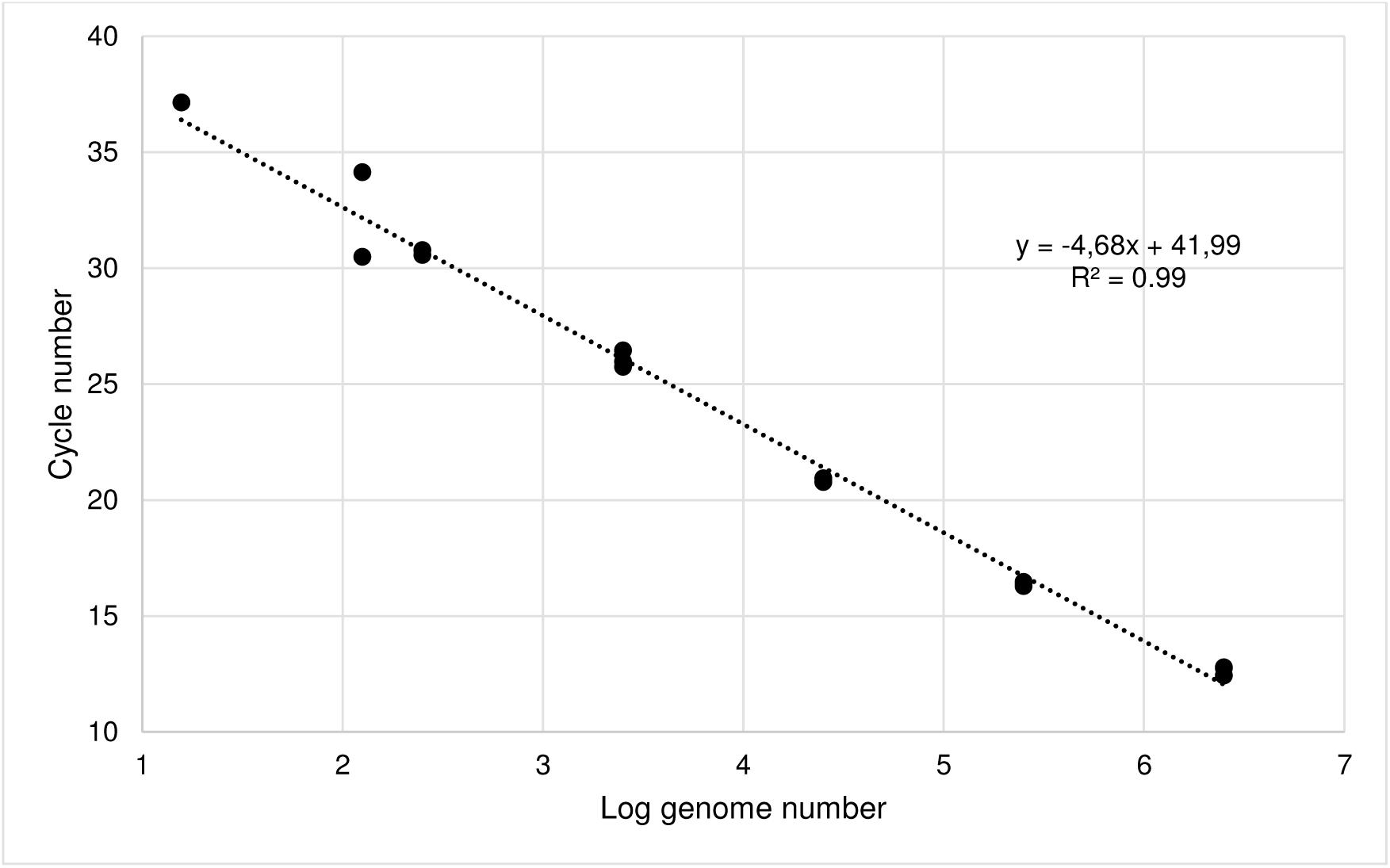
Standard curve established using real-time ZKIR PCR with serial dilutions of *K. pneumoniae* ATCC13883^T^ DNA from 7.5 ng to 45 fg.

### Analytical sensitivity of the ZKIR assay on soil samples

In order to assess the performance of the ZKIR assay for detection of Kp directly from soil samples, two soils (A and V) were spiked with bacterial concentrations ranging from 1.5 × 10^-1^ cfu/g to 1.5 × 10^4^ cfu/g. When soil samples were enriched in LB for 24 h and processed as described above, Kp was detected in all spiked microcosms except in two out of the 3 microcosms of soil A inoculated with the lowest Kp concentration. When the soil/LB suspension was tested prior to incubation, Kp was not detected in any of the spiked microcosms of soil V, whereas in soil A, Kp could only be detected at the highest concentration (1.5 × 10^4^ cfu/g). Finally, when the ZKIR assay was performed using purified metagenomic DNA from soil, positive results were only observed in microcosms spiked with 1.5 × 10^4^ and 1.5 × 10^3^ cfu/g in both soils, whereas no positive signal was observed at lower concentrations. The enrichment step thus appeared critical to reach high sensitivity.

### Comparison of the ZKIR real-time PCR and culture-based methods for the detection of Kp in environmental samples

After enrichment and ZKIR real-time PCR, Kp was detected in 41 out of 96 (42.7 %) assayed environmental samples, when the 1:10 dilution was used as template DNA (Table 1). When the 1:100 dilution was used, Kp was detected in a lower number of positive samples (38/96). The Ct values were in the range 18.4 - 36.2. The 96 samples were processed in parallel with the culture-based method. Kp was not detected in any of the ZKIR-negative samples: when presumptive colonies were detected, they were always identified by MALDI-TOF MS as non-Kp, and belonged to closely related species such as *K. oxytoca*, *Raoultella* spp. and *Serratia* spp. In contrast, Kp was isolated in 37 out of the 41 ZKIR positive samples. These isolates were all identified as either *K. pneumoniae* or *K. variicola* by MALDI-TOF MS.

Genomic characterization of 31 of these isolates (1 was contaminated and 5 were not yet available) revealed a dominance of Kp1 (n= 20, 65%), followed by Kp3 (n=10, 32%) and Kp4 (n=1, 3%; Table S3). Population diversity analysis based on MLST (39) and cgMLST (23) revealed a high genotypic diversity, with 23 STs and 25 cgMLST types. In five cases, the strain detected in the soil was the same as the one present in the leaves and/or roots (*e.g.*, SB6439 and SB6440; Table S3), showing colonization of several plant sites by the same strain. Only 2 isolates belonging to STs commonly found in the clinical settings were detected (ST37 and ST76). Interestingly, from the predicted O-types, the O3-type represented 50% of Kp population, which contrasts with the nosocomial situation, where types O1 and O2 are dominant (2, 40), but not with human carriage where the O3-type is also dominant (O3-31%, O1-19% and O2-17%; our unpublished results). Also contrasting with the clinical epidemiology of Kp, a low level of antibiotic resistance and virulence genes was observed, with 93.5% of the strains presenting an ancestral ‘wild-type’ susceptibility genotype (Table S3). The number of strains harboring plasmids detected in these environmental samples was also low (n=9, 29%), as well as the number of plasmid-encoded heavy metal tolerance genes (mainly the silver and copper tolerance clusters). These results contrast with clinical and animal isolates, where plasmids and metal tolerance clusters are common (42, Rodrigues, unpublished results). As expected (1, 36), all Kp3 isolates harbored the *nif* cluster responsible for nitrogen fixation. Interestingly, the *nif* cluster was also present in one Kp1 isolate from soil (SB6181). The phylogenetic analysis (**Figure S2**) of the *nif* cluster from our environmental isolates compared to a panel of reference strains (36) revealed that strain SB6181 (Kp1) branched within *K. variicola* strains (Kp3), showing that the *nif* cluster in this Kp1 strain was acquired via horizontal gene transfer from a *K. variicola* donor. Kp3 was also the inferred donor of the *nif* gene cluster for Kp5 and Kp6 *nif*-positive strains (Figure S2).

## Discussion

Routes of transmission of ecologically generalist human pathogens are usually complex and poorly understood. Proper risk management requires a holistic approach which has been theorised as the One Health concept (42). Despite the fact that the number of human infections caused by members of the Kp complex are on the rise and are increasingly resistant to antimicrobial treatment (1, 43), the ecology of Kp remains poorly understood. Identification of the various habitats in which Kp strives, and the routes of transmission to humans, for example through specific types of food, is critical in order to limit exposure. In this prospect, large-scale sampling is needed to define the sources of Kp contamination and such surveys will require sensitive, reliable and cost-efficient detection methods.

Several molecular assays for the detection and/or identification of *Klebsiella* from clinical or food and environmental samples have been previously proposed. These methods allow detecting mainly *K. pneumoniae* (*sensu stricto*) and *K. variicola* (Figure S1). However, the Kp complex currently encompasses 7 taxa. Our *in-silico* analyses showed that some previously described PCR assays could be useful for the identification of phylogenetic subsets of the Kp complex (Kp150233a for Kp1, *bla*_OKP_ for Kp2/Kp4 and *bla*_LEN_ for Kp3/Kp5).

Given that all members of the Kp complex can cause infections in humans or animals, developing a novel method with the ability to detect the Kp complex by targeting exhaustively all currently known Kp phylogroups would be an important advance. Although several previous targets were designed for the entire Kp complex, they predate recent taxonomic advances and have therefore not been validated on the entire phylogenetic breadth of this bacterial group. Here, we found by *in-silico* approaches that *khe* and *tyrB* assays lack specificity, which may negatively affect Kp detection efforts especially when testing microbiologically complex samples from the environment, such as soil.

We therefore aimed to develop a novel real-time PCR assay, and used an intergenic region located adjacent to the previously proposed target gene *khe*. Using well-defined reference strains representative of current taxonomy (20, 31, 32), we show that the novel ZKIR PCR real-time assay allows the accurate detection of all members of the Kp complex. We found complete specificity and sensitivity (*i.e*., phylogenetic coverage) of this assay based on our reference strains panel. To improve analytical sensitivity, a 24 h enrichment culture followed by an easy, fast and cost-effective sample processing was implemented, upstream of molecular detection using the ZKIR assay. This was necessary for complex matrices such as soil or sewage, where microbial diversity and abundance are high, in which case direct detection of one particular group of organisms, which might be present only at low abundance, is challenging. This procedure turned out to be sensitive enough to detect 1 single Kp bacterium in 5 g of soil. The ZKIR assay also appeared slightly more sensitive than the SCAI culture-based method, itself previously shown to be highly sensitive and to recover most Kp members (35, 44). Therefore, this fast and easy novel molecular method represents a powerful approach for screening large numbers of samples. This will spare the time-consuming handling of numerous presumptive colonies necessary for confirmation of their identification, given that *K. oxytoca*, *Raoultella, Serratia* and *Enterobacter* are able to form colonies on SCAI agar, with morphological characteristics similar to Kp (35, 45). As such, the ZKIR protocol was significantly faster than culture, as results were available on average 26 h after sampling (24 h enrichment, sample treatment, PCR) while up to 96 h were necessary when using the culture-based method (24 h enrichment, plate incubation, colony purification, MALDI-TOF identification).

Interestingly, the implementation of the ZKIR PCR real-time assay evidenced a high (43%) detection rate of Kp from environmental samples, which represent niches that are still underexplored for Kp presence and biological characteristics. The high detection rate was largely confirmed by culture, suggesting a low rate of false positive qPCR results. This novel and original sampling strategy allowed us to explore the genomic features of environmental Kp populations. A strong contrast with Kp isolates typically recovered in the clinical setting was observed. First, a high genetic diversity was observed, with no predominant sublineage. This situation contrasts with MDR or hypervirulent Kp populations, from which frequent sublineages (so-called high-risk clones) are recovered. Second, environmental Kp isolates were almost devoid of antibiotic resistance and virulence genes, in conspicuous contrast with clinical samples (46). Finally, the genomic characteristics of environmental Kp isolates revealed interesting biological features, such as a frequent O3 O-antigen type and the horizontal transfer of nitrogen fixation gene cluster into Kp1, the main phylogroup associated with human infections. These data call for further studies into the biology of soil Kp isolates, which may reveal interesting novel adaptive strategies of this important generalist pathogen. Finally, our findings suggest that environmental Kp populations differ from clinical Kp populations, which implies indirect and possibly complex epidemiological links between environmental and clinical Kp. The ZKIR PCR real-time assay developed herein is expected to enable future large-scale studies into this important question.

In conclusion, the ZKIR assay is a new tool for Kp detection that is highly specific, sensitive, reliable and cost-effective. To accelerate uptake of this method, the corresponding protocol was released publicly on protocols.io (<u>dx.doi.org/10.17504/protocols.io.7n6hmhe</u>). This simple method can easily be implemented in laboratories equipped with real-time PCR thermocyclers (MedVetKlebs consortium, unpublished results). Using the ZKIR method after a short culture enrichment step greatly enhances sensitivity. As this rapid screen of samples allows one to focus only on ZKIR positive samples for more labour-intensive downstream microbiological isolation and characterization, it is our hope that it will contribute to advance knowledge on the biology of environmental Kp and on the reservoirs and transmission routes of this increasingly important group of pathogens.

## SUPPLEMENTAL MATERIAL

**SUPPLEMENTAL FILE:** 4 PDF files, 1 Excel file

## ACKNOWLEDGEMENTS

We thank Alexis Criscuolo for access to an early version of the JolyTree software tool, Julien Guglielmini for providing us access to in-house scripts for mapping the target genes, and Juan Sebastian Lopez Fernandes for assistance with genomic analyses.

This work is part of the MedVetKlebs project, a component of the One Health European Joint Programme, and has thereby received funding from the European Union’s Horizon 2020 research and innovation programme under Grant Agreement No 773830.

